# Domestic goats can follow the direction of human voices to solve a hidden-object task

**DOI:** 10.64898/2025.12.08.692563

**Authors:** Stuart K Watson, Christian Nawroth, Alan G. McElligott, Federico Rossano, Simon W Townsend

**Affiliations:** Department of Evolutionary Anthropology, University of Zürich, Zürich, Switzerland; Research Institute for Farm Animal Biology, Dummerstorf, Germany; Department of Infectious Diseases and Public Health, Jockey Club College of Veterinary Medicine and Life Sciences, City University of Hong Kong, Hong Kong SAR; Centre for Animal Health and Welfare, Jockey Club College of Veterinary Medicine and Life Sciences, City University of Hong Kong, Hong Kong SAR; Department of Cognitive Science, University of California, San Diego, USA

**Keywords:** goats, reference, perception, domestication, acoustic processing

## Abstract

The capacity for animals to produce and comprehend vocalisations that provide referential acoustic cues to their eliciting cause (e.g. predator-specific alarm calls and food calls) is a highly adaptive means of maximising the benefits of group living, and has been widely studied in diverse species. However, an underexplored dimension of referentiality is the ability to process the direction in which a vocalisation is emitted as a cue towards its referent. Here, we replicated an experimental design previously applied to dogs, chimpanzees and human infants to investigate whether domestic goats (*Capra hircus*) can use human voices as a directional cue in a hidden-object task. Twenty-nine goats from a UK sanctuary participated in three experimental conditions. In each condition, goats were individually presented with a human experimenter obscured by a barrier, and two identical containers, one of which was baited with food. In the ‘reward directed speech’ condition, the experimenter vocalised excitedly towards the baited container while sitting closer to the un-baited container, and then the goat was able to select which container to explore. While substantial inter-individual variation existed, on average, subjects chose the baited container at above chance level across the four trials. Two control conditions explored alternative explanations for this result: a “no speech” condition, in which the experimenter was silent but remained in the same location as in the test condition, and a “non-reward directed speech” condition, in which the experimenter directed their voice away from both containers. Subjects showed no evidence of choosing the baited container at above chance level in either of these control conditions. We conclude that goats, like dogs, but not chimpanzees, are capable of attending to the directional cues provided by human voices, and discuss the possible role of domestication in the taxonomic distribution of this ability.

## Introduction

The ability for signal-receivers to infer the eliciting cause (i.e. ‘referent’) of a vocalisation based solely on its spectro-temporal features has received widespread attention in the last decades, both with regards to its ecological significance (Fichtel, 2020; Smith, 2017), as well as the light it can potentially shed on the evolution of semantic language (Fedurek & Slocombe, 2011; Townsend & Manser, 2013). Across various species, it has been demonstrated that an impressive degree of specificity can be inferred by receivers regarding the referents of these signals. This includes the type (Manser, 2013; Seyfarth & Cheney, 1986), location (Berthet et al., 2019) and/or relative urgency (Townsend et al., 2012) of a predator; or alternatively the presence (Evans & Marler, 1994), quality (Slocombe & Zuberbühler, 2005, 2006) and/or quantity (Di Bitetti, 2005) of a food resource. Interestingly, less research has focused on cues unrelated to the acoustic form of the signal itself, despite the fact that these can also encode referential information. For example, the direction in which a vocalisation is broadcast can allow receivers to infer the location of an external object in a three-dimensional environment, just as manual gestures, such as pointing, allow individuals to visually triangulate objects in space. This ability to use voice directionality as a referential cue would have obvious benefits in terms of reducing uncertainty regarding the precise location of a referent, and has been posited as a crucial faculty underlying joint-attention in human infants (Carpenter et al., 1998; Rossano et al., 2012a).

Initial work on the ability to process voice directionality as a referential cue has been carried out in only a few species, however this has been done using a standardised experimental approach, which is crucial for generating directly comparable data. Specifically, Rossano and colleagues experimentally examined the ability of human infants (12-16 months) (Rossano et al., 2012a), chimpanzees (*Pan troglodytes*) (Rossano et al., 2012a) and domestic dogs (*Canis familiaris*) (Rossano et al., 2014) to use the direction of an unseen human’s voice in an object-choice task. In this setup, the authors presented subjects with a visually obstructive barrier and a visible container at each end of the barrier (one of which was secretly baited with a reward). An experimenter concealed themselves from the subject by sitting behind the barrier, close to the empty container, and spoke excitedly in the direction of the baited container. Human infants and dogs, but not chimpanzees, directed their search towards the baited container significantly more often than expected by chance. Moreover, control conditions in which the experimenter was either silent or faced away from both containers while vocalising, ruled out the use of olfactory or other auditory cues to find the food. Even puppies (8-14 weeks) performed at above chance level, except for individuals who had limited exposure to humans, suggesting that there may be an additional learned component to the behaviour. The role of experience was further emphasised by a later study (Langner et al., 2023), which demonstrated that dogs perform better at this task when the speaker is their owner compared to an experimenter, and when the speech included relevant content (e.g. an attention-getter for the dog). Based on their findings and lack of positive results for chimpanzees, Rossano et al. (Rossano et al., 2014) posit that this skill may have developed in dogs as a result of domestication shaping their social cognition.

While intuitively compelling, the argument that dogs’ social cognition has been extensively shaped by the domestication process initiated over 10,000 years ago is a topic of considerable debate, with some contrasting evidence based on comparative work with wolves (Range & Marshall-Pescini, 2022b, 2022a). Furthermore, contrary to the notion that sensitivity to human voice direction is generally selected for during domestication, only one out of 16 domesticated piglets (*Sus scrofa domestica*) could locate a reward using human voice direction alone, while others needed the cue to be combined with pointing (Bensoussan et al., 2016). While this study had crucial methodological differences from the studies of Rossano et al. (Rossano et al., 2012a, 2014), this finding may suggest that either dogs’ ability to use human voice direction in a referential way is due to some pre-existing trait present in their ancestry unrelated to domestication. Alternatively, it could be that the domestication processes for companion animals and livestock do not necessarily give rise to similar socio-cognitive abilities. Expanding the taxonomic survey of the non-human animal ability to follow human voices as a referential cue will shed further light on this topic.

Goats (*Capra hircus*) are highly social ruminants that live in fission-fusion social groups with stable hierarchies (Briefer et al., 2015; Stanley & Dunbar, 2013) and were first domesticated approximately 10.5k years ago, making them one of humanity’s oldest domesticated species (MacHugh et al., 2017; MacHugh & Bradley, 2001). The socio-cognitive abilities of goats with regard to their sensitivity to human visual cues have, therefore, been a popular topic of research in recent years (Deutsch et al., 2025; Mason, Semple, et al., 2024; Nawroth, 2017). It has been found that goats prefer positive over negative human facial expressions (Nawroth et al., 2018), are sensitive to human gaze and head orientation (Kaminski et al., 2005; Nawroth et al., 2015, 2016; Nawroth & McElligott, 2017), and can even follow a variety of pointing gestures (Nawroth et al., 2020, 2023a). Meanwhile, in the acoustic domain, goats can discriminate the emotional valence of human voices (Mason, Semple, et al., 2024) and also detect when the voice and face of familiar humans are incongruent (Mason et al., 2025). However, whether they can use the directionality of voices as a referential cue has yet to be tested.

Here, we replicate the experimental setup of Rossano et al. (Rossano et al., 2012a, 2014) (previously used with infants, chimpanzees, and dogs) to generate directly comparable data on the ability of domestic goats to process the directionality of human vocalisations as a referential cue. As in Rossano et al. (Rossano et al., 2012a, 2014), we tested the goats in three conditions: i) a ‘reward directed speech’ condition, where the obscured experimenter vocalised excitedly towards the baited container, ii) a ‘no speech’ control condition, in which the experimenter did not vocalise at all, and iii) a ‘non-reward directed speech’ control condition, in which the experimenter vocalised excitedly in an orthogonal direction to both baited and empty containers so that it was not a useful directional cue. If goats are capable of processing directional acoustic cues from an unseen speaker without additional cues, we predicted that most individuals would select the baited container more often than chance level (50%). Conversely, success rates in each of the control conditions would not reliably differ from chance level unless goats were utilising confounding cues such as olfaction to find the food.

## Methods

### Study site and subjects

All data were collected at Buttercups Sanctuary for Goats, UK (https://www.buttercups.org.uk/) in September 2016, in Kent, U.K. (51°13′15.7″, N 0°33′05.1″E). The sanctuary was open to visitors and animals were provided with ad libitum hay, grass and water, and commercial concentrate relative to age and condition. At night they were housed in individual pens, or small groups (average pen size 3.5 m2). A total of 36 adult goats were tested, but seven were excluded from full participation (three for motivation reasons, three were tested in a pilot condition, and one was excluded due to experimenter error), leaving a final sample of 29 individuals (9 females, 20 males) who were tested on all three experimental conditions. All goats had prior experience as subjects in behavioural research and were highly habituated to human interaction (Nawroth et al., 2018, 2020). For handling, a leash was put on the collar that was released once the subject was supposed to make a choice. Subjects were used to this procedure for routine procedures and previous tests. None of the goats attempted to get rid of the leash and easily followed the human when they were moving.

### Apparatus

A large wooden barrier (150 x 200 cm) was placed in a testing enclosure (7m x 5.3m), equidistant from both sides of the enclosure (Figure 1). Two identical containers (red buckets) were placed at each end of the wooden barrier. All trials were recorded using digital video cameras (Sony HCRCV 190E Camcorder). When containers were baited with a food item, this was always a piece of uncooked pasta. Subjects had collars during the course of the experiment (and were habituated to this already).

**Figure 1.**
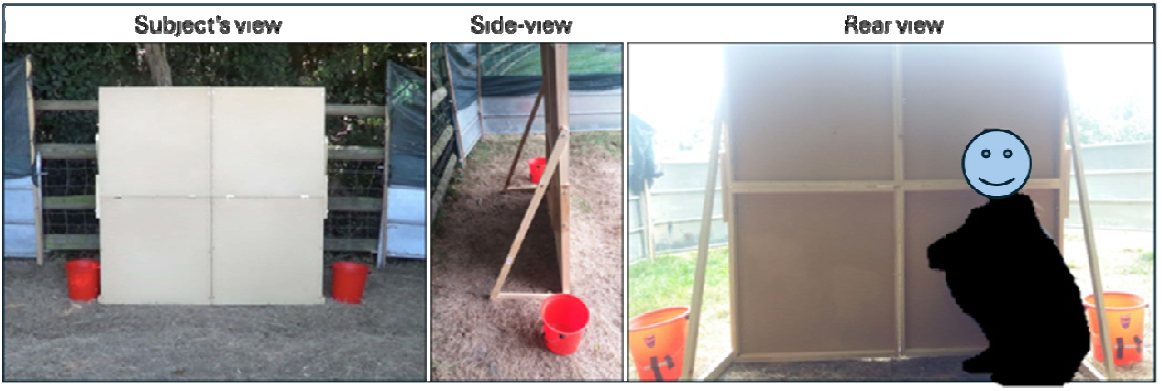
View of the experiment setup from the perspective of a subject, and behind the screen. Research assistant anonymised for preprint.

### Procedure

#### Pre-test familiarisation phase

Before each of the experimental conditions, a pretest familiarisation phase was conducted. In this, the test enclosure was set up just as it would be during the experimental phase (Figure 2), and the subject was led through the testing enclosure by Experimenter 2. The subject was able to explore the space behind the barrier to ensure it was fully habituated to the apparatus and setup. Experimenter 2 led the goat to the starting point to initiate the pretest. First, Experimenter 1 stood centrally behind the barrier. Appearing above the centre of the barrier in view of the subject, Experimenter 1 showed a piece of food and called the subject’s name. Then, Experimenter 1 visibly placed the food in one of the containers. The animal was able to track the food because Experimenter 1’s hand was always in full view of the subject. Which container was baited was pseudo-randomised, with no more than two consecutive trials on the same side. The test phase started when the goat successfully retrieved the food in two consecutive trials (one on each side) to demonstrate they understood the task.

**Figure 2.**
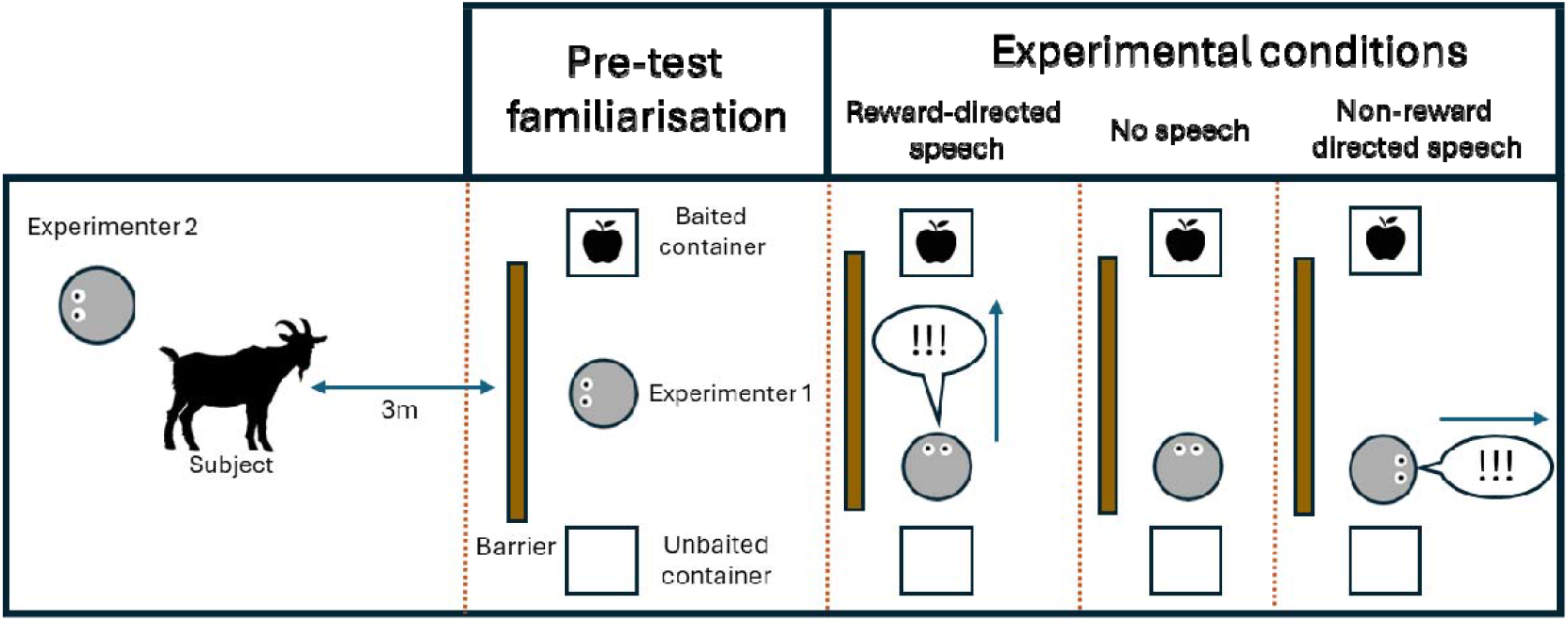
Schematic for experimental procedure for each condition. The location of the baited container was counterbalanced between trials.

#### Experimental phase

Three experimental conditions (‘reward directed speech’, ‘no speech’, and ‘non-reward directed speech’ conditions; Figure 2) were carried out to explore the ability of goats to use the direction of human speech as a cue for finding food. For each of these conditions, four consecutive trials were carried out for every subject, with placement of the baited container counterbalanced across trials. Therefore, each subject underwent a total of 12 experimental trials. For all individuals, the reward directed speech condition was run first, followed by the no speech and then non-reward directed speech conditions. All four trials from a single condition were carried out back-to-back within the same session (max. 8 minutes), and then the next condition was run several days later. All testing was carried out between 11am and 4pm.

#### ‘Reward directed speech’ condition

The subject was led by Experimenter 2 to the start position, oriented towards the barrier. Experimenter 2 also faced away from the barrier to avoid providing any cues. Experimenter 1 then stood behind the barrier and called the subject’s name. Then, ducking down behind the barrier, Experimenter 1 placed the food in one of the containers. Experimenter 1 then moved the containers from behind the barrier such that they were visible at each end of the barrier. Experimenter 1, who was concealed behind the barrier, oriented their face and body towards the baited container, but positioned themselves closer (<30cm apart) to the empty container. This ensured that the subject was not simply using the side of the barrier located closest to the voice to guide their choice. In this position, Experimenter 1 excitedly spoke a predefined sentence: “*ja guck mal, ja guck mal da, das ist ja fein!*” (“*Oh look, look there, this is great!*”) twice, and then remained silent. Experimenter 2 released the animal after two utterances, so they could choose one of the containers. If the subject moved more than 1.5m towards the baited container, it was rewarded with the food inside. If the animal approached the empty container, no food reward was received. After each trial, the animal was redirected by Experimenter 2 away from the barrier, and back to the testing position if another trial was to take place.

#### ‘No speech’ condition

A ‘no speech’ control condition was used to determine whether there were any non-voice related cues (e.g., olfaction) which the subjects might have used to locate the baited container. In this, the procedure was identical to that of the ‘reward directed speech’ condition, except that Experimenter 1 was completely silent (Figure 2) and the subject was released after 5s instead of after the first sentence. Four consecutive trials were carried out for every subject, with placement of the baited container counterbalanced across trials.

#### ‘Non-reward directed speech’ condition

To determine whether it was the actual direction of the voice that goats were using to guide their search for food, we ran a ‘non-reward directed speech’ control condition, which was identical to the ‘reward directed speech’ condition except that Experimenter 1 oriented their head orthogonally to the wooden barrier, so that their voice carried no directional cues to the baited container (Figure 2). Four consecutive trials were carried out for every subject, with placement of the baited container counterbalanced across trials.

### Video coding

For each trial, the behaviour of the subject was coded individually by two observers, one of whom was blind to the location of the baited container. A container was coded as chosen by an animal when it touched the container, or moved more than 1.5m towards it from the starting position. Inter-rater reliability between the two observers was almost perfect (Cohen’s κ = 0.994).

### Statistical analysis

We sought to explore i) whether subjects chose the correct container at above chance levels (50%) in each condition, and ii) if the probability of subjects’ success differed between the reward directed speech and condition and the control conditions. This was done by fitting a single Bayesian linear mixed effect model using the ‘bernoulli’ family and a logit link function. Whether an individual successfully chose the baited container in a given trial (yes/no) was fitted as a binary response measure, and experimental condition (‘reward directed speech’, ‘no speech’, and ‘non-reward directed speech’ conditions) was fitted as a fixed effect. Random effects were fitted for ‘subject ID’ and ‘trial number’ to control for non-independence of data. All models used weakly regularising priors to minimise the risk of overfitting. Evidence of an effect of a predictor was determined by the extent to which the 89% credible interval for the posterior estimate of a predictor crossed zero, with coefficients that did not cross zero indicating evidence of a robust effect. The model was run using the package ‘brms’ (Bürkner, 2017) in R (R Core Team, 2019) and RStudio (RStudio Team & others, 2015) with two chains of 10,000 iterations each. For all models, we assessed model chain convergence by inspecting trace plots, rhat values (all equal to 1.00) and effective sample sizes for each parameter (all well over 1000). Data and scripts used for analysis can be found at the following repository: https://tinyurl.com/goatdirection

### Ethical note

The study was approved by the Animal Welfare and Ethical Review Board committee of Queen Mary University of London (Ref. QMULAWERB072016). The subjects were housed at Buttercups Goat Sanctuary, a registered UK charity who rescue goats from diverse backgrounds, who also granted permission for the work to be carried out. Animal care and all experimental procedures were in accordance with the 2016 ASAB/ABS guidelines for the use of animals in research (Association for the Study of Animal Behaviour, 2016). Individual separation from conspecifics, as performed in this experiment, is a common husbandry practice and did not cause any visible stress to the animals. If any signs of stress had been shown (e.g. loud vocalisations, restless wandering/escape attempts, and rejection of food rewards), the experiment would have ceased immediately. Written informed consent was obtained from individuals depicted in potentially identifiable images included in this article, which were taken by the authors.

## Results

The mean success rate for choosing the baited container was as follows for each condition (Figure 3, Figure 4 for individual trials): reward directed speech (60%), non-reward directed speech (49%) and no speech (47%). Our binomial GLMM indicated that goats had an 89.8% posterior probability (calculated from log-transformed model outputs, Table 1) of performing at above chance level (set at 50% success) across the four trials in the reward directed speech condition, while the average estimated probability of success on any given trial was 59.6% (89% CI: 51.0–68.1%). Conversely, there was only a 43.5% posterior probability of performing above chance level across the four trials in the no speech condition (Posterior probability of success for any given trial: 49.1%, 89% CI: 40.4% - 57.9%), and 24.7% for the non-reward directed speech condition (Posterior probability of success for any given trial: 46.5%, 89% CI: 37.9% - 55.0%). Furthermore, the model estimated a 93.8% posterior probability that subjects performed better in the reward directed speech condition than in the no speech condition, and a 97.3% probability compared to the non-reward directed speech condition (Table 1). In summary, goats generally performed at above chance level in the reward directed speech condition, and reliably better than in either control condition, which did not elicit performance above chance.

**Table 1.**
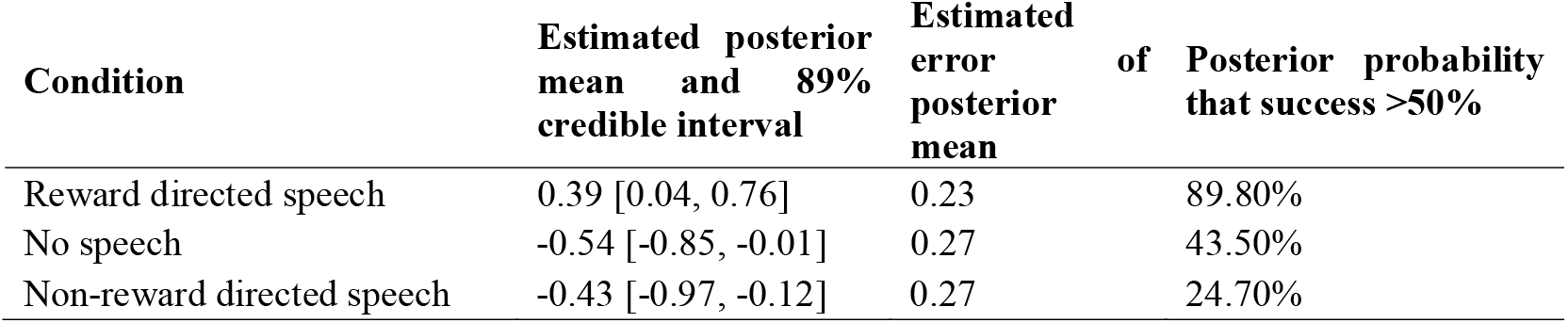
Model outputs per experimental condition. The reward directed speech condition was set as the intercept, therefore, estimates for ‘non-reward directed speech’ and ‘no speech’ conditions are reported relative to this. Credible intervals which do not cross zero can be taken as robust evidence of an effect. Therefore, the reward directed speech condition elicited successful container choice at above chance level, and reliably more often than both control conditions. Posterior probability that success was greater than 50% was calculated using an inverse log transformation of the corresponding posterior mean.

**Figure 3.**
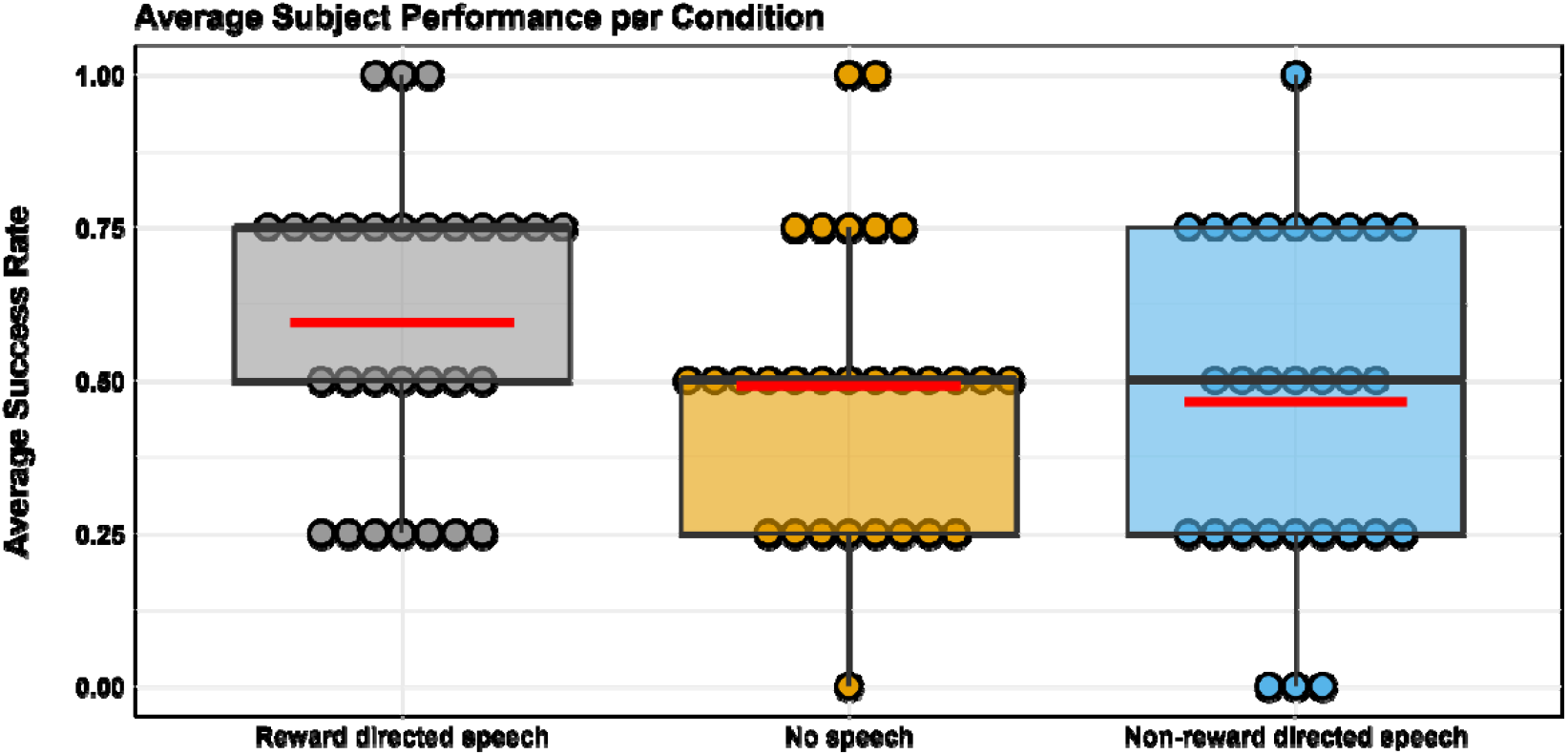
Summary of median (thick grey line) and mean (red line) success rates per condition, where each dot represents the success rate (out of four trials) for a single subject. Each subject was tested in all three conditions. See Figure 4 for individual performance across conditions.

**Figure 4.**
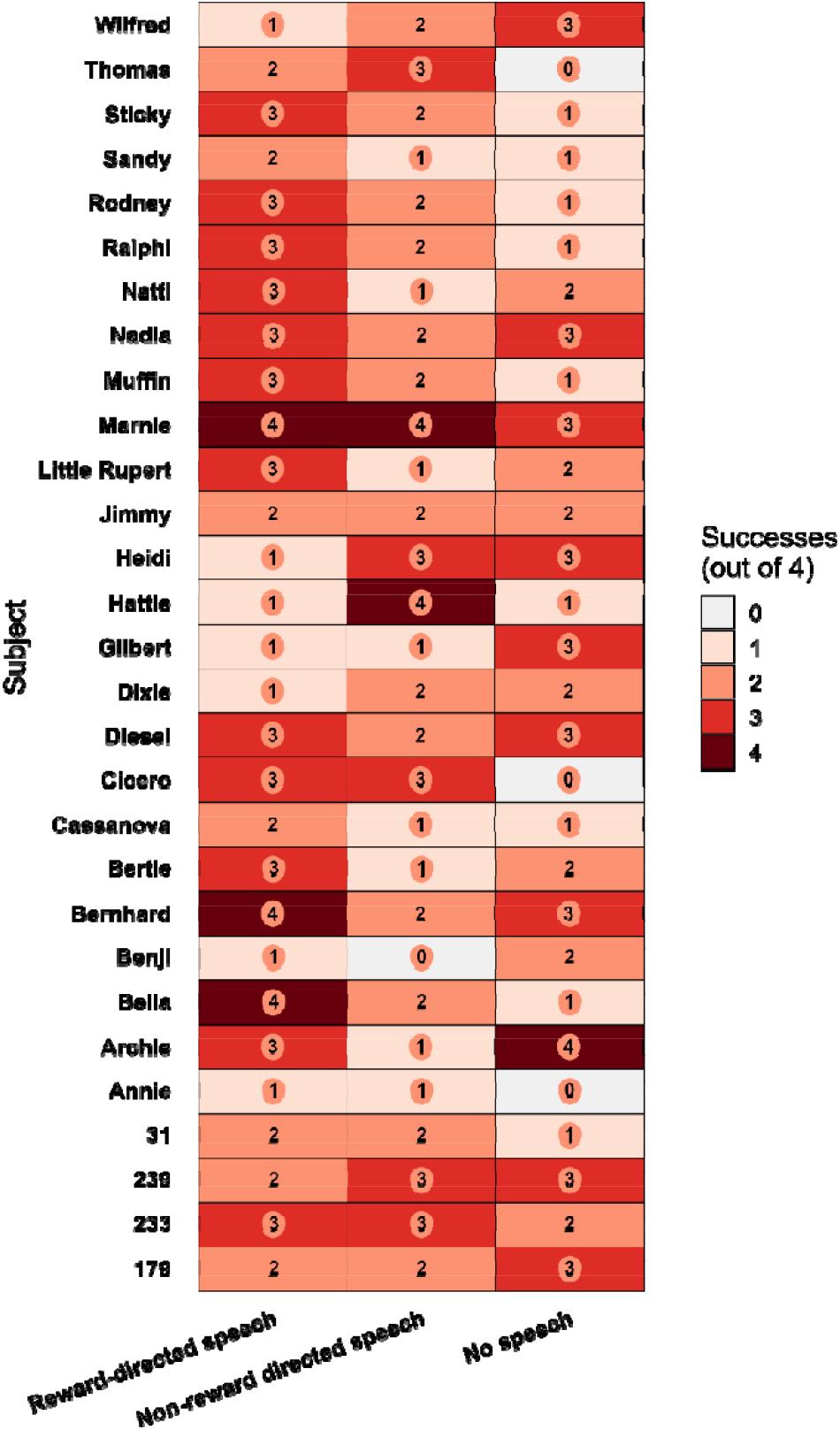
Individual number of successes out of four trials for each experimental condition.

## Discussion

The ability to follow the directionality of vocalisations may be an important cue in decoding the referent of a signal (Rossano et al., 2012a). The ability to process human voices in this manner has previously been established as present in the majority of dogs tested (Rossano et al., 2014), but absent in most chimpanzees (Rossano et al., 2012a), suggesting it has potentially been shaped by domestication. To expand the taxonomic survey of this ability to further domesticated species, we tested the ability of domestic goats to follow human voice direction. We found that goats are capable of following the direction of an unseen human’s voice, as our subjects were generally able to find hidden food at above chance level based solely on this cue. Crucially, control conditions in which the voice was either directed away from both baited and unbaited containers, or where there was no voice at all, confirmed that this result could not be explained by attention to any other visual, acoustic or olfactory cues. This ability to process the directionality of heterospecific vocalisations adds to the growing body of literature demonstrating the complexity of social cognition used by goats for navigating relationships with humans (Mason, Briefer, et al., 2024; Nawroth, 2017).

The capacity to use human voice directionality in a referential manner highlights an intriguing degree of cognitive flexibility in goats. The processing of referential signals by non-human species is generally studied in terms of receivers having an explicit association between an external stimulus and the acoustic form of a vocalisation, typically demonstrated by playback experiments causing receivers to behave as though the stimulus were present even when it was not (Magrath et al., 2015; Townsend & Manser, 2013). Here, the referent of the signal had to be flexibly decoded according to the dynamic cue of human voice directionality. This capacity arguably allows receivers to achieve a level of referential specificity in signal processing that would otherwise be impossible in a system lacking a detailed semantic component (e.g., “the container to my left”). Applying this ability to heterospecific signals belies a further layer of flexibility on the part of receivers (Magrath et al., 2015). Exploring whether goats also process the directionality of conspecific signals in this way might shed further light on whether this trait is specific to human relationships (emerging via domestication and/or ontogenetically through our sample’s extensive experience with humans) or represents a more general capacity that can be applied across communicative contexts.

It has long been argued that the impressive sensitivity of dogs to the social and communicative cues of humans is a product of domestication, but agreement on this is not unanimous (Range & Marshall-Pescini, 2022b, p. 202, 2022a), and data from other domesticated species is crucial to drawing firm conclusions. Interestingly, dogs and goats had descriptively very similar rates of success in this particular task (∼60% in goats vs. ∼63% in dogs; Rossano et al., 2014), while one-year-old humans (Rossano et al., 2012b) performed higher at around an 80% success rate. Meanwhile, chimpanzees which were also tested by Rossano et al. (2012) did not succeed at above chance level. Taken at face value, it is tempting to posit that these convergent lines of evidence support the view that the selective pressures of domestication have shaped the socio-cognitive abilities of both dogs and goats. However, a recent direct experimental comparison between the abilities of domestic and wild goats to interpret very salient human social and physical cues did not find any robust differences between these samples in both domains (Nawroth et al., 2023b), indicating that at least part of this package of traits may have been present in their non-domesticated ancestor. Whether the more subtle ability to follow human vocal directionality is among these ancestral traits would need to be established with further comparative research. Currently, at least ∼1.2 billion domestic goats exist globally, predominantly as livestock (FAOStats, 2024). Therefore, in addition to the scientific value of this research program, advancing our understanding and appreciation of the behaviour and cognition of goats, particularly with regards to their relationship with humans, may have important downstream impacts on public perception and welfare for this species (Bastian et al., 2012; Cox & Gaston, 2015; Mason, Briefer, et al., 2024; Stokes, 2007; Sweet et al., 2023; Zobel & Nawroth, 2020).

While most goats were successful in the reward directed condition, substantial inter-individual variation was evident, with almost half of the sample performing at or below chance in the experimental condition (Figure 3, Figure 4). This suggests that even with the latent ability to track the direction of human speech, achieving accuracy is challenging and prone to error, and likely to be impacted by differences in motivation and/or skill. Many genetic, developmental and personality-related factors (Briefer et al., 2015; Nawroth et al., 2017) could drive individual differences of this nature, but the details of subjects’ exposure to humans is one particularly plausible factor. For example, in Rossano et al. (2014), puppies who had lived as pets with humans performed at above chance level in the task, while individuals with less human exposure did not. Furthermore, the only chimpanzee within Rossano et al.’s (2012) sample of 16 individuals to perform at above chance level (10 successes out of 12 trials) had been raised in a human home during their early life. Similarly, the domesticated piglets tested by Bensoussan et al. (Bensoussan et al., 2016), who failed to solve a similar task, had just two weeks of habituation to humans before testing. Given that our sample of goats lived in a sanctuary and came from diverse backgrounds, it seems likely that the nature and extent of individual goats’ relationship with humans may have driven similar phenotypic plasticity in their sensitivity to human vocal cues (Webster & Rutz, 2020). Supporting the importance human exposure on how social cognition is deployed, even non-domesticated kangaroos solicit human assistance during an unsolvable experimental task, which the authors of the study attribute to the sample’s previous positive experiences with humans (McElligott et al., 2020). Future work could explore this possibility in goats by comparing data from individuals with extensive, versus relatively little, direct human interaction.

A noteworthy detail of the present study is that our subjects spontaneously (without prior training) made the relevant associations between the experimenter’s speech, its direction, and the goat’s prior understanding that one of the containers was baited. Our data therefore demonstrate not only the inherent capability of goats to process the directionality of human speech, but also to spontaneously infer the implications of this cue within a novel context. It is indeed possible that some subjects did, in fact, follow the direction of the speech, but failed to associate the vocal cues with the details of the object-choice. This could be explored in the future through more passive visual/acoustic mismatch experiments, taking measures of surprise such as gaze duration or nasal temperature (Ermatinger et al., 2019) from subjects who observe an individual facing in one direction, and then hear an acoustic cue travelling in either the same or opposite direction. It would also be worthwhile to explore how combinations of visual and acoustic cues are processed to improve inferential accuracy, and how they are weighed against one another when they conflict (Scandurra et al., 2018, 2020). Finally, our experiment used a German sentence as a stimulus since this was the native language of the experimenter, but it is possible that the goats would have been more readily attuned to the task if we had used a more familiar English phrase. However, while this aspect of the design may have constrained the overall success rate of our goats, it arguably also reinforces the robustness of our findings since subjects were generally able to accurately infer the referent of the signal even in an unfamiliar language.

In conclusion, goats are readily able and spontaneously motivated to use the directionality of human speech to infer the presence and location of food. This data adds to the accumulating body of literature examining this largely unexplored aspect of referential processing, and provides further evidence that goats are highly sensitive to human social cues (Nawroth et al., 2015, 2016, 2020). While the exact role of domestication in the emergence of this trait is unclear (Nawroth et al., 2023b; Range & Marshall-Pescini, 2022b, 2022a), further insight can be provided by broadening the taxonomic survey of these abilities to yet further domesticated and non-domesticated species.

## Acknowledgements

We thank Lea Wagner for her assistance in collecting the data. We thank all staff and volunteers at Buttercups Sanctuary for Goats (http://www.buttercups.org.uk) for their excellent help, expertise and access to the animals. The authors have no competing interests to declare.

## Data availability

All data and R scripts used for analysis can be accessed at the following repository: https://tinyurl.com/goatdirection

## References

Association for the Study of Animal Behaviour. (2016). Guidelines for the treatment of animals in behavioural research and teaching. Animal Behaviour, 111, i–ix. 10.1016/S0003-3472(15)00461-3

Bastian, B., Loughnan, S., Haslam, N., & Radke, H. R. M. (2012). Don’t Mind Meat? The Denial of Mind to Animals Used for Human Consumption. Personality and Social Psychology Bulletin, 38(2), 247–256. 10.1177/0146167211424291

Bensoussan, S., Cornil, M., Meunier-Salaün, M.-C., & Tallet, C. (2016). Piglets Learn to Use Combined Human-Given Visual and Auditory Signals to Find a Hidden Reward in an Object Choice Task. PLOS ONE, 11(10), e0164988. 10.1371/journal.pone.0164988

Berthet, M., Mesbahi, G., Pajot, A., Cäsar, C., Neumann, C., & Zuberbühler, K. (2019). Titi monkeys combine alarm calls to create probabilistic meaning. Science Advances, 5(5), eaav3991. 10.1126/sciadv.aav3991

Briefer, E. F., Oxley, J. A., & McElligott, A. G. (2015). Autonomic nervous system reactivity in a free-ranging mammal: Effects of dominance rank and personality. Animal Behaviour, 110, 121–132. 10.1016/j.anbehav.2015.09.022

Bürkner, P.-C. (2017). brms: An R Package for Bayesian Multilevel Models Using Stan. Journal of Statistical Software, 80(1). 10.18637/jss.v080.i01

Carpenter, M., Nagell, K., Tomasello, M., Butterworth, G., & Moore, C. (1998). Social Cognition, Joint Attention, and Communicative Competence from 9 to 15 Months of Age. Monographs of the Society for Research in Child Development, 63(4), i. 10.2307/1166214

Cox, D. T. C., & Gaston, K. J. (2015). Likeability of Garden Birds: Importance of Species Knowledge & Richness in Connecting People to Nature. PLOS ONE, 10(11), e0141505. 10.1371/journal.pone.0141505

Deutsch, J., Langbein, J., & Nawroth, C. (2025). Learning and cognition in goats. In Small Ruminant Welfare, Production and Sustainability (pp. 291–309). Academic Press. 10.1016/B978-0-443-22201-6.00012-8

Di Bitetti, M. S. (2005). Food-associated calls and audience effects in tufted capuchin monkeys, Cebus apella nigritus. Animal Behaviour, 69(4), 911–919. 10.1016/j.anbehav.2004.05.021

Ermatinger, F. A., Brügger, R. K., & Burkart, J. M. (2019). The use of infrared thermography to investigate emotions in common marmosets. Physiology & Behavior, 211, 112672. 10.1016/j.physbeh.2019.112672

Evans, C. S., & Marler, P. (1994). Food calling and audience effects in male chickens, Gallus gallus: Their relationships to food availability, courtship and social facilitation. Animal Behaviour, 47(5), 1159–1170. 10.1006/anbe.1994.1154

FAOStats. (2024). Crops and livestock products. https://www.fao.org/faostat/en/#data/QCL

Fedurek, P., & Slocombe, K. E. (2011). Primate Vocal Communication: A Useful Tool for Understanding Human Speech and Language Evolution? Human Biology, 83(2), 153–173. 10.3378/027.083.0202

Fichtel, C. (2020). Monkey Alarm Calling: It Ain’t all Referential, or is It? Animal Behavior and Cognition, 7(2), 101–107. 10.26451/abc.07.02.04.2020

Kaminski, J., Riedel, J., Call, J., & Tomasello, M. (2005). Domestic goats, Capra hircus, follow gaze direction and use social cues in an object choice task. Animal Behaviour, 69(1), 11–18. 10.1016/j.anbehav.2004.05.008

Langner, L., Žakelj, S., Bolló, H., Topál, J., & Kis, A. (2023). The influence of voice familiarity and linguistic content on dogs’ ability to follow human voice direction. Scientific Reports, 13(1), 16137. 10.1038/s41598-023-42584-2

MacHugh, D. E., & Bradley, D. G. (2001). Livestock genetic origins: Goats buck the trend. Proceedings of the National Academy of Sciences, 98(10), 5382–5384. 10.1073/pnas.111163198

MacHugh, D. E., Larson, G., & Orlando, L. (2017). Taming the Past: Ancient DNA and the Study of Animal Domestication. Annual Review of Animal Biosciences, 5(Volume 5, 2017), 329–351. 10.1146/annurev-animal-022516-022747

Magrath, R. D., Haff, T. M., Fallow, P. M., & Radford, A. N. (2015). Eavesdropping on heterospecific alarm calls: From mechanisms to consequences: Interspecific eavesdropping. Biological Reviews, 90(2), 560–586. 10.1111/brv.12122

Manser, M. B. (2013). Semantic communication in vervet monkeys and other animals. Animal Behaviour, 86(3), 491–496. 10.1016/j.anbehav.2013.07.006

Mason, M. A., Briefer, E. F., Semple, S., & McElligott, A. G. (2024). Goat Emotions, Cognition, and Personality. In S. Mattiello & M. Battini (Eds.), The Welfare of Goats (pp. 77–120). Springer Nature Switzerland. 10.1007/978-3-031-62182-6_3

Mason, M. A., Semple, S., Marshall, H. H., & McElligott, A. G. (2024). Goats discriminate emotional valence in the human voice. Animal Behaviour, 209, 227–240. 10.1016/j.anbehav.2023.12.008

Mason, M. A., Semple, S., Marshall, H. H., & McElligott, A. G. (2025). Do goats recognise humans cross-modally? PeerJ, 13, e18786. 10.7717/peerj.18786

McElligott, A. G., O’Keeffe, K. H., & Green, A. C. (2020). Kangaroos display gazing and gaze alternations during an unsolvable problem task. Biology Letters, 16(12), 20200607. 10.1098/rsbl.2020.0607

Nawroth, C. (2017). Invited review: Socio-cognitive capacities of goats and their impact on human– animal interactions. Small Ruminant Research, 150, 70–75. 10.1016/j.smallrumres.2017.03.005

Nawroth, C., Albuquerque, N., Savalli, C., Single, M.-S., & McElligott, A. G. (2018). Goats prefer positive human emotional facial expressions. Royal Society Open Science, 5(8), 180491. 10.1098/rsos.180491

Nawroth, C., Martin, Z. M., & McElligott, A. G. (2020). Goats Follow Human Pointing Gestures in an Object Choice Task. Frontiers in Psychology, 11. 10.3389/fpsyg.2020.00915

Nawroth, C., & McElligott, A. G. (2017). Human head orientation and eye visibility as indicators of attention for goats (Capra hircus). PeerJ, 5, e3073. 10.7717/peerj.3073

Nawroth, C., Prentice, P. M., & McElligott, A. G. (2017). Individual personality differences in goats predict their performance in visual learning and non-associative cognitive tasks. Behavioural Processes, 134, 43–53. 10.1016/j.beproc.2016.08.001

Nawroth, C., von Borell, E., & Langbein, J. (2015). ‘Goats that stare at men’: Dwarf goats alter their behaviour in response to human head orientation, but do not spontaneously use head direction as a cue in a food-related context. Animal Cognition, 18(1), 65–73. 10.1007/s10071-014-0777-5

Nawroth, C., von Borell, E., & Langbein, J. (2016). ‘Goats that stare at men’—revisited: Do dwarf goats alter their behaviour in response to eye visibility and head direction of a human? Animal Cognition, 19(3), 667–672. 10.1007/s10071-016-0957-6

Nawroth, C., Wiesmann, K., Schlup, P., Keil, N., & Langbein, J. (2023a). Domestication and breeding objective did not shape the interpretation of physical and social cues in goats (Capra hircus). Scientific Reports, 13(1), 19098. 10.1038/s41598-023-46373-9

Nawroth, C., Wiesmann, K., Schlup, P., Keil, N., & Langbein, J. (2023b). Domestication and breeding objective did not shape the interpretation of physical and social cues in goats (Capra hircus). Scientific Reports, 13(1), 19098. 10.1038/s41598-023-46373-9

R Core Team. (2019). R: A language and environment for statistical computing. [Computer software]. R Foundation for Statistical Computing. https://www.R-project.org/

Range, F., & Marshall-Pescini, S. (2022a). Comparing wolves and dogs: Current status and implications for human ‘self-domestication.’ Trends in Cognitive Sciences, 26(4), 337–349. 10.1016/j.tics.2022.01.003

Range, F., & Marshall-Pescini, S. (2022b). Domestication Hypotheses Relating to Behaviour and Cognition: Which Are Supported by the Current Data? In F. Range & S. Marshall-Pescini (Eds.), Wolves and Dogs: Between Myth and Science (pp. 335–373). Springer International Publishing. 10.1007/978-3-030-98411-3_11

Rossano, F., Carpenter, M., & Tomasello, M. (2012a). One-Year-Old Infants Follow Others’ Voice Direction. Psychological Science, 23(11), 1298–1302. 10.1177/0956797612450032

Rossano, F., Carpenter, M., & Tomasello, M. (2012b). One-Year-Old Infants Follow Others’ Voice Direction. Psychological Science, 23(11), 1298–1302. 10.1177/0956797612450032

Rossano, F., Nitzschner, M., & Tomasello, M. (2014). Domestic dogs and puppies can use human voice direction referentially. Proceedings of the Royal Society B: Biological Sciences, 281(1785), 20133201. 10.1098/rspb.2013.3201

RStudio Team & others. (2015). RStudio: Integrated development for R [Computer software]. RStudio, Inc., Boston, MA. http://www.rstudio.com

Scandurra, A., Alterisio, A., Aria, M., Vernese, R., & D’Aniello, B. (2018). Should I fetch one or the other? A study on dogs on the object choice in the bimodal contrasting paradigm. Animal Cognition, 21(1), 119–126. 10.1007/s10071-017-1145-z

Scandurra, A., Pinelli, C., Fierro, B., Di Cosmo, A., & D’Aniello, B. (2020). Multimodal signaling in the visuo-acoustic mismatch paradigm: Similarities between dogs and children in the communicative approach. Animal Cognition, 23(5), 833–841. 10.1007/s10071-020-01398-9

Seyfarth, R. M., & Cheney, D. L. (1986). Vocal development in vervet monkeys. Animal Behaviour, 34(6), 1640–1658. 10.1016/S0003-3472(86)80252-4

Slocombe, K. E., & Zuberbühler, K. (2005). Functionally Referential Communication in a Chimpanzee. Current Biology, 15(19), 1779–1784. 10.1016/j.cub.2005.08.068

Slocombe, K. E., & Zuberbühler, K. (2006). Food-associated calls in chimpanzees: Responses to food types or food preferences? Animal Behaviour, 72(5), 989–999. 10.1016/j.anbehav.2006.01.030

Smith, C. L. (2017). Referential signalling in birds: The past, present and future. Animal Behaviour, 124, 315–323. 10.1016/j.anbehav.2016.08.015

Stanley, C. R., & Dunbar, R. I. M. (2013). Consistent social structure and optimal clique size revealed by social network analysis of feral goats, Capra hircus. Animal Behaviour, 85(4), 771–779. 10.1016/j.anbehav.2013.01.020

Stokes, D. L. (2007). Things We Like: Human Preferences among Similar Organisms and Implications for Conservation. Human Ecology, 35(3), 361–369. 10.1007/s10745-006-9056-7

Sweet, F. S. T., Noack, P., Hauck, T. E., & Weisser, W. W. (2023). The Relationship between Knowing and Liking for 91 Urban Animal Species among Students. Animals, 13(3), 488. 10.3390/ani13030488

Townsend, S. W., & Manser, M. B. (2013). Functionally Referential Communication in Mammals: The Past, Present and the Future. Ethology, 119(1), 1–11. 10.1111/eth.12015

Townsend, S. W., Rasmussen, M., Clutton-Brock, T. H., & Manser, M. B. (2012). Flexible alarm calling in meerkats: The role of the social environment and predation urgency. Behavioral Ecology, 23(6), 1360–1364. 10.1093/beheco/ars129

Webster, M. M., & Rutz, C. (2020). How STRANGE are your study animals? Nature, 582(7812), 337–340. 10.1038/d41586-020-01751-5

Zobel, G., & Nawroth, C. (2020). Current state of knowledge on the cognitive capacities of goats and its potential to inform species-specific enrichment. Small Ruminant Research, 192, 106208. 10.1016/j.smallrumres.2020.106208

